# Echolocation calls of some bat species in western Uganda

**DOI:** 10.64898/2025.12.12.694018

**Authors:** Laura N. Kloepper, Reese N. Fry, Olivia Maliszewski, Robert S. Hahn, James A. Simmons, Andrea Megela Simmons

## Abstract

With the rise of accessible recording technology, passive acoustic monitoring can be an affordable and rapid way to assess species richness, even when individual animals cannot be captured due to regulatory or practical obstacles. Motivated by the relative lack of data and in partnership with the local populace, we recorded echolocation calls of freely-flying bats across six locations in rural western Uganda using opportunistic passive acoustic recordings.

Frequency-modulated echolocation calls were recorded at all six locations, while constant-frequency calls were recorded only at sites near entrances to caves. Preliminary species identifications were made using Kaleidoscope Pro, habitat distribution maps for Uganda, and by reference to published work. We make our acoustic recordings publicly available to serve as a resource for further explorations of the richness of bat species in Uganda.

## Introduction

Bats (order Chiroptera) inhabit a wide range of ecological niches worldwide and are particularly abundant in the tropics (Denzinger and Schnitzler 2013). Because they occupy a high trophic level, have slow reproductive rates, and function as pollinators and in insect control, bats serve as crucial bioindicators of ecological health (Jones et al. 2009; Stahlschmidt and Brühl 2012; Russo et al. 2021). Like many mammals, bats are threatened by anthropogenic activities including habitat loss, hunting, and climate change, and a large swath of the globe is considered data-deficient for bats (Frick et al. 2019). Sampling more bat species in broader parts of the world is of utmost importance to fully understand global patterns of bat biodiversity and population status, to develop improved methods for conservation, and to expand educational outreach to nonscientists (Krause 1993; Meyer et al. 2010; Barlow et al. 2015; López-Bosch et al. 2024).

In the past two decades, passive acoustic monitoring has become increasingly popular as a technique for monitoring the distribution and population size of vocal animals worldwide (Sueur et al. 2008; Sugai et al. 2019). Because echolocating bats emit species-specific calls during foraging and navigation, acoustic monitoring offers an effective, low-cost, low-invasive way to estimate the size, health, and biodiversity of bat populations, particularly in understudied parts of the world (e.g., Taylor et al. 2005; Jennings et al. 2008; Andreassen et al. 2014; Kloepper et al. 2016; Rydell et al. 2017; Gibb et al. 2019; Howard et al. 2022; Hending et al. 2022; Grunwald et al. 2024; Roemer et al. 2025). One of the requirements for acoustic identification is the need for call recording and reference libraries that have been verified by species identification (via DNA, morphology, etc.) that can then be used to generate automatic call classifiers and to identify species in real time in the field. But, the availability of reference libraries and automatic classifiers is a limitation for ecologists desiring to conduct reliable species identification in many areas of the world. For example, the Kaleidoscope Pro software (Wildlife Acoustics, Maynard, MA, USA), a popular commercial option for auto-identification, has species classifiers for broad regional categories (North America, Europe, South America, Neotropics, South Africa) representing only 7-8% of the approximately 1,500 known species of bat. An open-access global call library “ChiroVox” (Görföl et al. 2022) contains reference calls from 242 species spanning fourteen countries, but including only one African country (Liberia). Another limitation of automatic classifiers and call reference libraries is that the algorithms used for developing them are trained on stereotyped recordings from known individuals, either while resting, held in the hand, or flying alone in open environments. These classifiers have variable accuracy for less stereotyped calls, those from understudied species, or those recorded under realistic field conditions where signal-to-noise ratios are not optimal and when multiple conspecific and heterospecific bats are present (Jennings et al. 2008; Russo and Voigt 2016; Rydell et al. 2017; Brinkley et al. 2021; Roemer et al. 2021). Thus, even now, automatic classification must be ground-truthed against manual classification.

The continent of Africa contains great biological richness, with approximately 12 families and 300 species of bats identified to date (ACR 2024; Monadjem et al. 2024). As of 2009, approximately 95 bat species have been identified in Uganda alone (Thorn and Peterhans 2009). Echolocation calls of some of these species have been recorded from resting or freely-flying individual bats, and examined in the context of taxonomy, systematics, and biogeographical variables (Csorba et al. 2003; Monadjem et al. 2011, 2020, 2024; ACR 2024). There are, however, few online databases granting free access to echolocation call recordings of these species for further study and educational use.

To expand the potential of bat acoustic identification worldwide, scientists should increase the number, availability, and accessibility of recordings. Even when field conditions and permitting restrictions prohibit direct species identification from individual captures, acoustic recordings alone have value. They can be archived and analyzed retroactively once reference libraries have been developed for a region, providing insights into trends in bat biodiversity and distribution. Our work aims to contribute to the study of the richness of echolocating bat species in Uganda using opportunistic passive acoustic monitoring. As part of informal “bat walks” with the local populace, we recorded echolocation calls from free-flying bats at six different locations in rural southwestern Uganda. From these echolocation calls, we calculated frequency and temporal parameters and quantified acoustic entropy, a metric used to estimate acoustic diversity between habitats (Sueur et al. 2008). We suggest some preliminary species identifications based on the output of an automated classifier for South African bats (Brinkley et al. 2021; Kaleidoscope Pro), by reference to habitat distribution maps for Uganda (Monadjem et al. 2024), and by consulting the published literature (Kityo and Kerbis 1996; Taylor 1999; Csorba et al. 2003; Taylor et al. 2005; Monadjem et al. 2011, 2017, 2020; Webala et al. 2019; Grunwald et al. 2024; ACR 2024). We make our acoustic recordings available to contribute not only to increased understanding of bat richness and distribution in Uganda, but also to the growing collection of acoustic datasets for global bat biodiversity monitoring.

## Materials and methods

### Recording locations

Opportunistic acoustic recordings were conducted in April-May 2015 across six recording sites in rural western Uganda: Murchison Falls (MUR), Mutolere (MUT), Mweya (MWE), Lake Nyabikere (NYA), Rubuguri (RUB), and Ruhanga (RUH). These sites were identified in partnership with local inhabitants who were asked where bats in the area might be observed, and who then accompanied the researchers to observe and discuss the field recordings.

Bat calls were recorded at a sampling rate of 192 kHz (the limits of the recording device) using a single Dodotronic Momimic (Dodotronic, Castel Gandolfo, Italy) microphone connected to a SoundDevices 744T recorder (Sound Devices, Reedsburg, WI, USA). The recorder was placed in one position at each site, with the microphone either held in hand pointing towards the direction of the bats emerging from a cave (MUT, RUB, RUH) or positioned on a flat surface oriented towards the sky as bats flew overhead (MUR, MWE, NYA). Recording heights and distance to the flying bats were not consistent across sites. Recordings were made opportunistically when bats were observed in each area, with recordings starting at sunset and lasting from 30 min up to 2 hours, depending on the amount of bat activity on the recording night. Due to permitting restrictions, bats were not captured for physical species identification.

### Call analysis and classification

Echolocating bats are broadly divided into two groups based on the major acoustic features of their echolocation calls, although bats actively vary these features during pursuit of prey (the pursuit sequence; Simmons et al. 1979). Constant frequency (CF-FM) bats are high duty-cycle echolocators, producing narrowband pulses with long durations relative to the signal period (Fenton et al. 2012). CF pulses can have several harmonics and peak frequencies up to 160 kHz. In some species, a short, frequency-modulated (FM) sweep occurs at the beginning and/or at the end of the CF component. As these bats approach prey, they adjust their calls so that the CF component decreases in duration and the FM component becomes more prominent (Simmons et al. 1979; Fenton et al. 2012). FM bats are low duty-cycle echolocators, emitting short downward-sweeping pulses separated by inter-pulse intervals that are longer than the duration of the emitted pulses. FM echolocation calls have multiple harmonics that decrease in frequency over a species-typical range from about 130 kHz down to 10 kHz. FM bats also actively alter their echolocation pulses as they approach an insect prey, such that the slope of the FM sweep becomes sharper and its duration shortens from ∼<10 ms in the search phase to ∼1 ms (feeding buzz) at the time of capture (Simmons et al. 1979). Even within these broad groups, there is considerable diversity in the structure of the echolocation call related to geographic location, microhabitat, foraging strategies, and the presence of conspecifics or other clutter (Obrist 1995; Denzinger and Schnitzler 2013). We focused on analyzing echolocation calls in the search phase of pursuit.

We analyzed the recorded audio files using both manual and automated analyses. Audio files were uploaded to Adobe Audition v. 2021 and converted to 16-bit WAV files. To reduce electrical and wind noise, files were processed through Adobe Audition’s noise and hiss reduction algorithms. Using MATLAB (v. 2021b; MathWorks, Natick, MA, USA), the processed audio files at each site were high-pass filtered (*highpass*; minimum order FIR filter with a stopband attenuation of 60 dB) at 10 kHz and then separated into one-minute-long file segments. From these files, we automatically extracted calls and characterized their spectral and temporal features using a custom energy detector written in MATLAB. This was accomplished by calculating the power spectrogram of the filtered audio between the cutoff frequency and the Nyquist frequency using the function *pspectrum* with the input argument ‘spectrogram’ and a time resolution of 5 ms. The algorithm identifies individual calls in the filtered files using the functions *findpeaks* and *envelope*. Onset and offset thresholds of an identified call are set as the maximum of.001 and 1/10th of the maximum amplitude of the upper envelope. The duration of the call is calculated using the nearest envelope-threshold crossing points to the maximum amplitude. The minimum call duration was set to 3 ms to eliminate feeding buzzes signaling successful prey capture. Thus, calls in the search and approach phases of insect pursuit (Simmons et al. 1979) are emphasized while feeding buzzes are disregarded. The algorithm also calculated entropy (spectral entropy, temporal entropy, entropy index) using formulas in Sueur et al. (2008).

We then compared the output of the automated algorithm to manual parameter extraction using Raven Pro 1.6 software (Cornell Lab of Ornithology, Ithaca, NY, USA) and/or Adobe Audition. Comparison of automated with manual classification (“ground-truthing”) is a standard practice in bioacoustic analyses, due to the limitations of automated detection (Rydell et al. 2017; Brinkley et al. 2021). Spectrograms of WAV files were computed at a sampling rate of 192 kHz and FFT size 65536. Using selection tables and power spectra, we extracted frequency (low frequency, high frequency, bandwidth) and temporal (duration) parameters. Pulse interval was not calculated because this parameter varies considerably across the pursuit sequence (Simmons et al. 1979) and in the presence of clutter (including from other bats; Obrist 1995; Surlykke and Moss 2000); moreover, the presence of multiple individuals with similar calls in our recordings complicated these measurements.

For tentative species identifications, we processed the audio files using Kaleidoscope Pro’s auto-classifier for South African bats (Brinkley et al. 2021). An auto-classifier for Ugandan bats is not available, and South Africa is the closest geographic match. The Kaleidoscope algorithm detects and classifies calls with reference to a known library based on call shape. It also provides information on the frequency and duration of detected calls. These classifications were then cross-referenced with previous literature on echolocation calls and species distributions of Ugandan bats (Kityo and Kerbis 1996; Csorba et al. 2003; Monadjem et al. 2011, 2020, 2024; ACR 2024).

For illustrations, spectrograms were computed in Adobe Audition 2021 and adjusted to reduce noise and hiss. Images were then imported to Adobe Photoshop 2025 to adjust color, brightness, and contrast.

## Results

We made a total of 470 min of audio recordings across the six locations, resulting in the following sample sizes of one-minute audio files: MUR=80, MUT=40, MWE=130, NYA=120, RUB=60, RUH=40. Our algorithm extracted the following total number of events at each location: MUR=12,430; MUT=7,784; MWE=5,594; NYA=13,660; RUB=9,237; RUH=6,936.

Assuming all of these detected events are bat calls (rather than, for example, insect or atmospheric sounds), then bat vocal activity at these six recording sites is high, ranging from 43-195 calls/min.

### Measures of entropy

Spectral entropy (H = 48.57, df = 5, p < 0.001), temporal entropy (H = 86.41, df = 5, p < 0.001), and entropy index (H = 45.85, df = 5, p < 0.001) differed among the recording locations (Fig. 1). We found spectral entropy varied significantly across most sites (Fig. 1a); spectral entropy was similar at MUR and NYA, and also similar between MUT, MWE, and RUH. On the other hand, both temporal entropy and entropy index remained relatively consistent across sites (Fig. 1b, c).

**Fig. 1.**
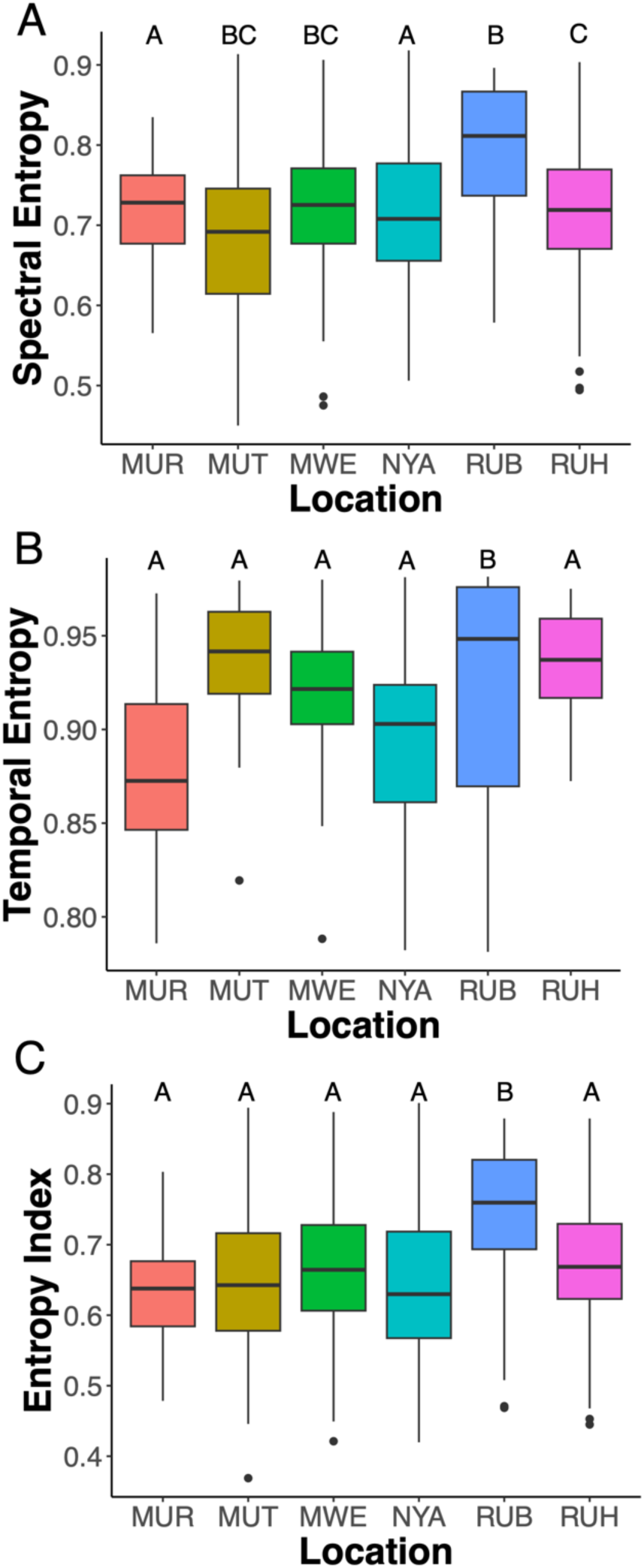
Boxplots of acoustic biodiversity measurements at the different recording locations. (A) spectral entropy, a measure of complexity in the frequency domain; (B) temporal entropy, a measure of complexity in the time domain; and (C) entropy index, the product of spectral and temporal energy and a measure of acoustic complexity. Solid black bars represent the median, shaded regions represent the inter-quartile range, vertical lines represent the minimum and maximums of the data, without outliers, and the circles represent outliers in the data. Letters indicate significantly different (p < 0.05) groups based on Dunn’s pairwise comparisons, with locations sharing the same letter(s) not statistically different (p > 0.05) from each other.

An exception was RUB, which showed higher values of spectral entropy, temporal entropy, and the entropy index than the other recording locations.

### Acoustic characteristics of echolocation calls

Visual inspection of spectrograms indicated that multiple individuals are present at each location; however, we did not attempt to count numbers of individual bats. At some sites, the variety of echolocation calls indicated the presence of multiple species. Below we summarize the main features of echolocation calls (CF-FM and FM) and make tentative species identifications.

### CF-FM calls

CF-FM calls were recorded at three sites (MUT, RUB, RUH), where recordings were made near cave entrances. Figure 2 shows examples of echolocation calls in the form of sample spectrograms from these three locations. At all three locations, calls have two prominent frequency bands (harmonics), one at around 44 kHz and the other at around 88 kHz. The stronger first harmonic is due to the impact of atmospheric attenuation (Lawrence and Simmons, 1982). The CF component of each harmonic is preceded and followed by a sharp FM sweep.

**Fig. 2.**
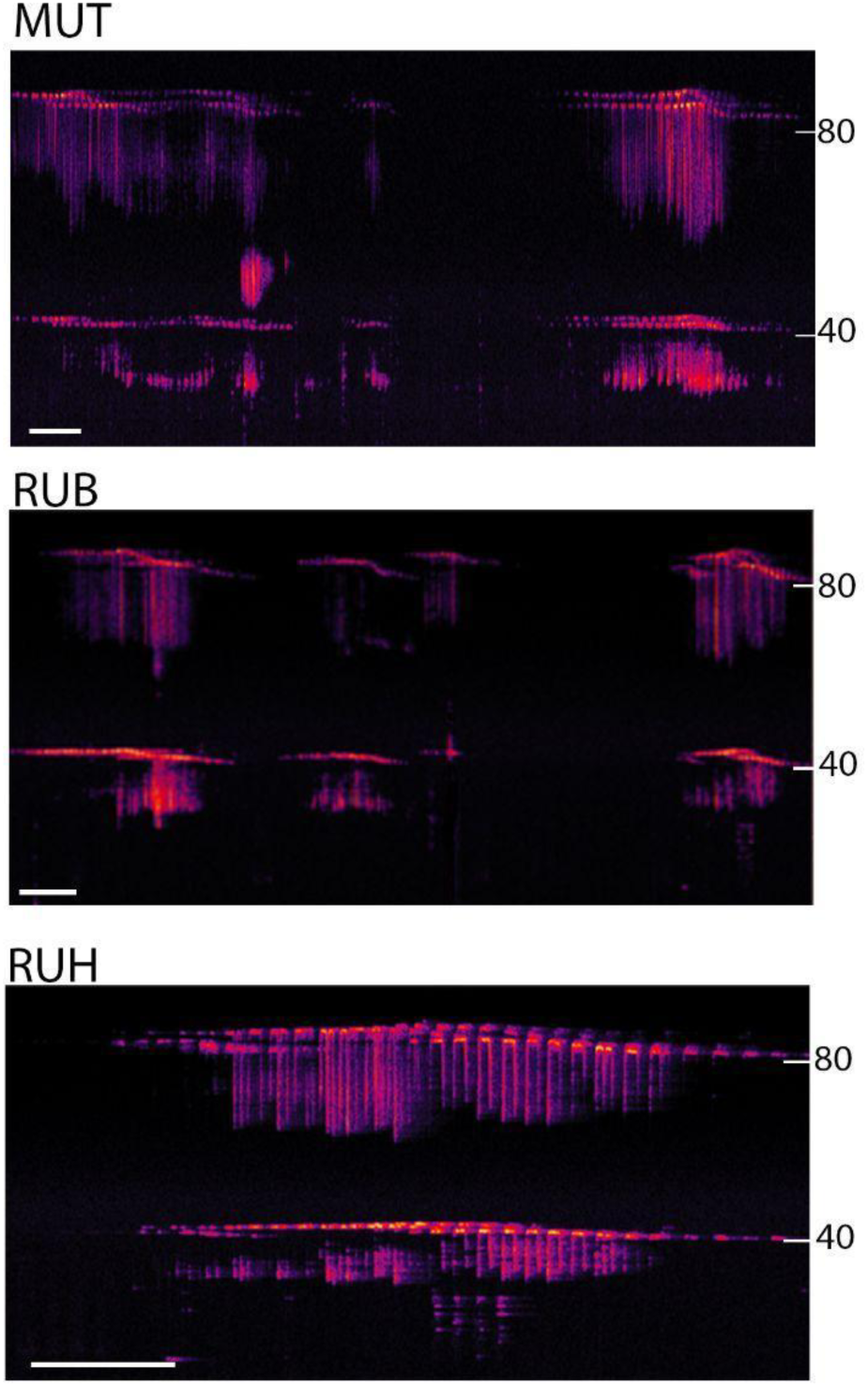
Sample spectrograms of CF-FM echolocation calls from three recording locations (MUT, RUB, RUH). The right y-axis of each spectrogram shows frequency in kHz. The x-axis of each spectrogram shows time, with the white scale bar indicating 500 ms. Spectrograms were computed in Adobe Audition (spectral frequency display, sample rate 192 kHz, 16 bit, FFT size 65536, Hamming window).

Multiple individuals are present at each site, and their calls often overlap. Table 1 gives mean values of harmonic frequencies and duration measured from non-overlapping calls.

**Table 1.** Acoustic features of individual CF-FM echolocation calls at three recording sites.

| Location | N | Peak frequency<br>1<sup>st</sup> harmonic (kHz) | Peak frequency<br>2<sup>nd</sup> harmonic (kHz) | CF duration<br>(ms) |
| --- | --- | --- | --- | --- |
| Mutolere | 21 | 43.65 ± .22 | 87.79 ± .51 | 43 ± 12 |
| Ruhanga | 22 | 44.03 ± .59 | 87.04 ± 1.0 | 36 ± 16 |
| Rubuguri | 22 | 43.37 ± .64 | 85.49 ± .94 | 40 ± 15 |

Values are means and standard deviations from individual calls (N) at each recording site

Because we were unable to capture any bats, we cannot make definite species determinations. The frequencies of the CF harmonics suggest that these calls are consistent with those recorded from rhinolophids. We used the database and distribution models introduced by Monadjem et al. (2024) to estimate the occurrence of Rhinolophus species with known echolocation calls at our recording sites. Fig. 3 illustrates a distribution map of *Rhinolophus acrotis*, with the approximate locations of our three recording sites superimposed. *R. acrotis* has been classified as a lineage of *R. clivosus* (Uvizl et al. 2024). The echolocation calls of *R. clivosus* have a resting frequency of ∼ 90 kHz, depending on recording location (Taylor 1999; Monadjem et al. 2017, 2020). We could find no published information on the echolocation calls of *R. acrotis*. Pye and Roberts (1970; Roberts 1972) recorded echolocation calls of another rhinolophid, *R. alcyone* (Halcyon horseshoe bat), captured in Uganda (Fig. 3), although not at our recording sites. The echolocation calls of this bat have a resting frequency of ∼87 kHz, within the range of our results.

**Fig. 3.**
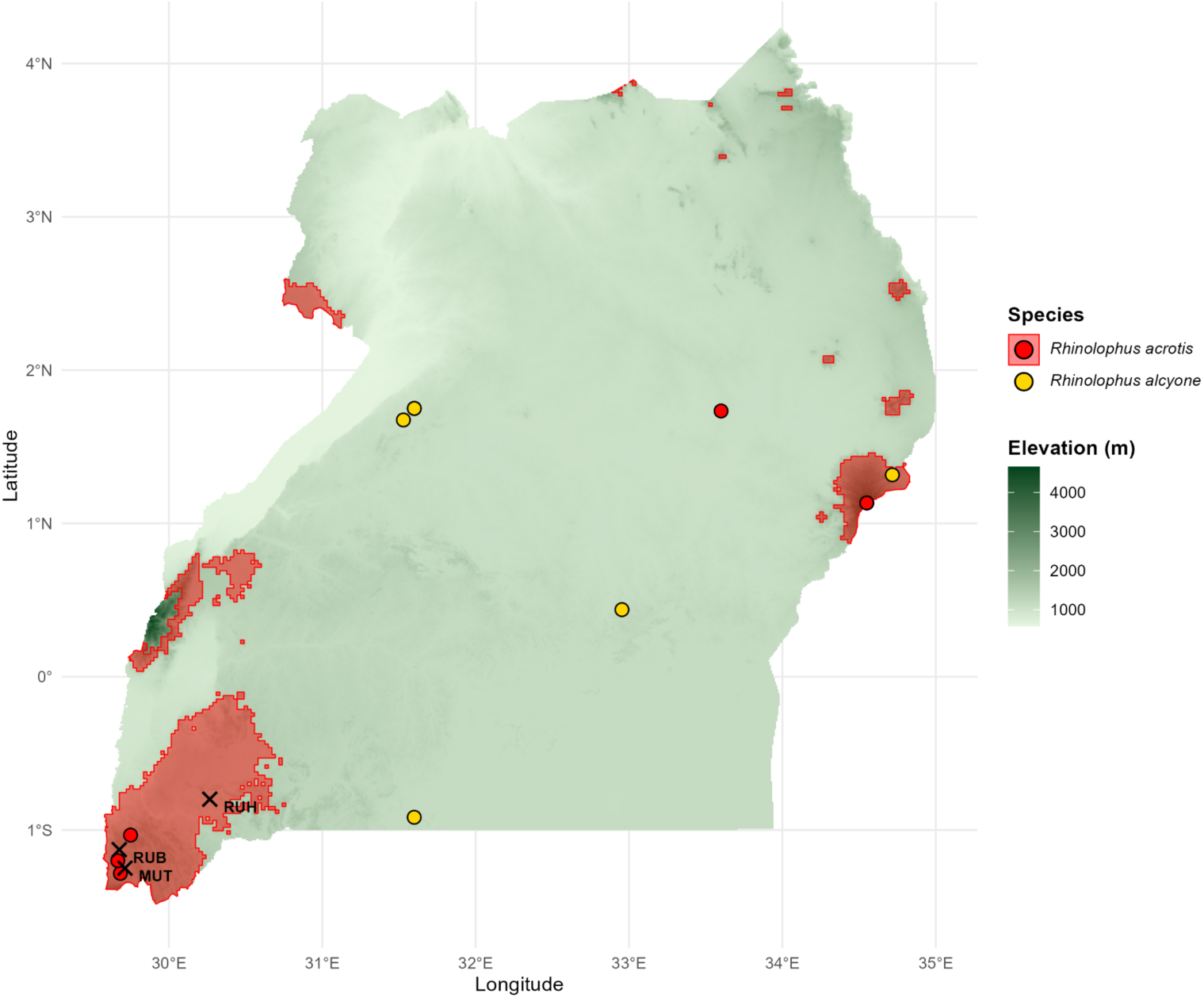
Output of the habitat model of Monadjem et al. (2024) showing the distribution of *R. acrotis* and of *R. alcyone* and the recording sites (MUT, RUB, RUH, X symbols) at which we observed CF-FM bats. Latitude is shown on the y axis and longitude on the x axis. Elevation is depicted in shades of green (right insert). The red outlines show suitable habitat for *R. acrotis* and the white circles show areas where this species has been captured. The yellow circles show areas where *R. alycone* has been captured; Monadjem et al. (2024) do not present data on suitable habitats for this species.

### FM calls

FM calls were prominent at three recording sites (MUR, MWE, NYA). At each of these sites, spectrogram visualization suggests that multiple species were present (Fig. 4).

**Fig. 4.**
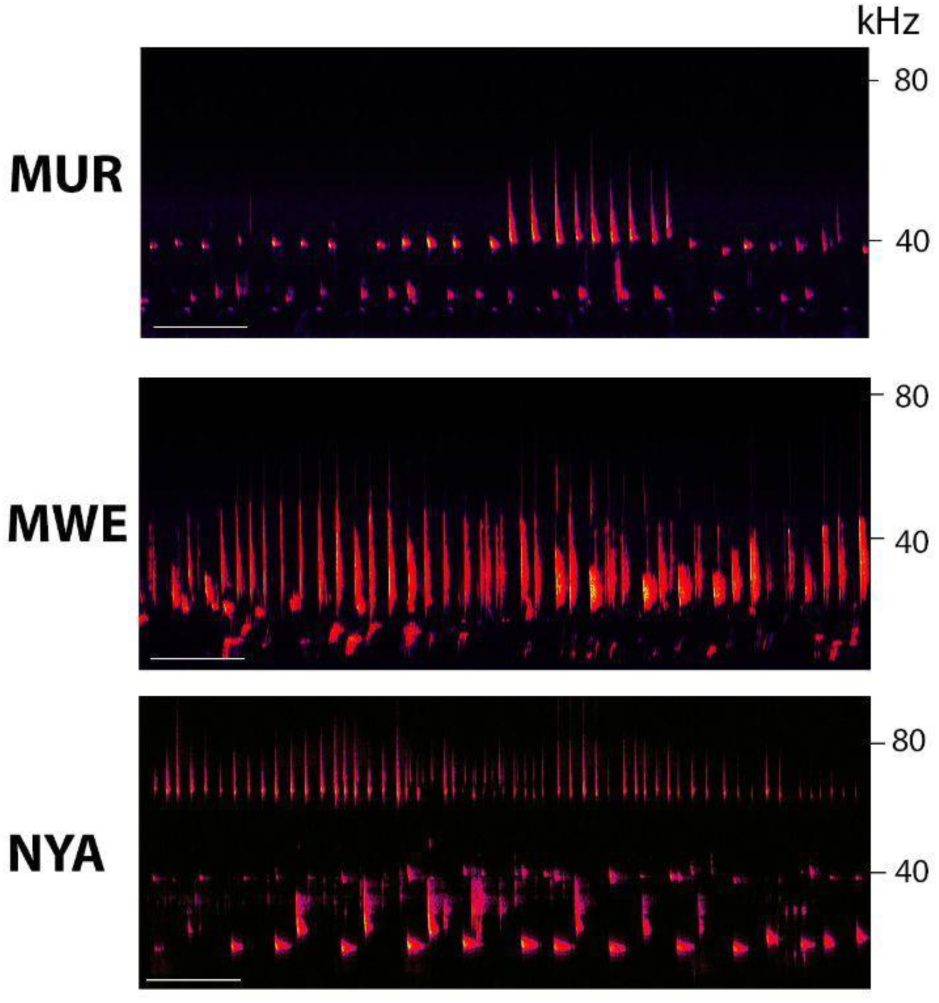
Sample spectrograms of FM echolocation calls at three recording sites (MUR, MWE, NYA). Multiple individuals and species are present at each site. The right y-axis of each spectrogram shows frequency in kHz. The x-axis shows time, with the white scale bar indicating 500 ms. Spectrograms were computed in Adobe Audition (spectral frequency display, sample rate 192 kHz, 16 bit, FFT size 65536, Hamming window). Durations of individual calls were on average 5.3 ms (range 3-9 ms) at MUR, 5.2 ms at MWE (range 3-10 ms), and 5.2 ms at NYA (range 3-11 ms).

Frequency characteristics of FM calls (high frequency distribution, low frequency distribution, bandwidth) at the three recording sites are plotted in Fig. 5. There is clear overlap in the high frequency energy (∼25-55 kHz) at these three sites. Some FM calls recorded at NYA (Lake Nyakibere) contained higher frequency energy between ∼60-100 kHz. The distribution of low frequency (first harmonic) energy shows a similar trend: overlap at three sites, with higher low frequency energy at NYA. These data suggest that at least two bat species were present at NYA.

**Fig 5.**
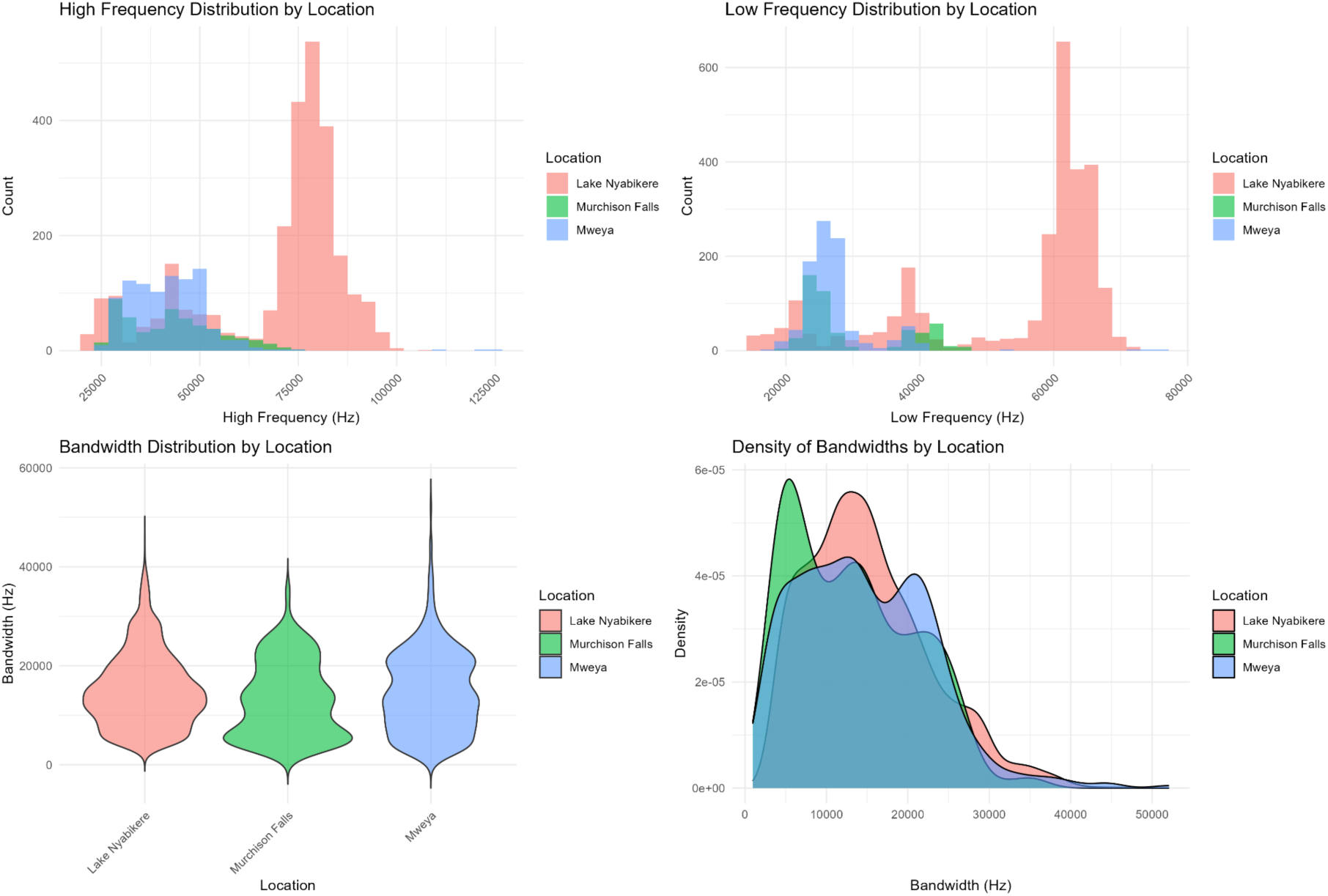
Distributions of frequency and bandwidth at sites at which FM echolocation calls were predominant (Lake Nyabikere, Murchison Falls, Mweya). Data were obtained by manual analysis in Raven Pro and in Adobe Audition, with cross-validation by the output of our MATLAB script. Distributions were calculated and plotted in RStudio Team (2020).

The vespertilionid *Afronycteris (Neoromicia) nana* (banana serotine bat) was documented by Monadjem et al. (2017, 2020, Swaziland and South Africa) and by Taylor-Boyd et al. (2025; Zambia) to echolocate within the frequency range of ∼ 67-∼86 kHz, consistent with the measurements we show in Fig. 6. We suggest that this bat was present at Lake Nyabikere.

**Fig. 6.**
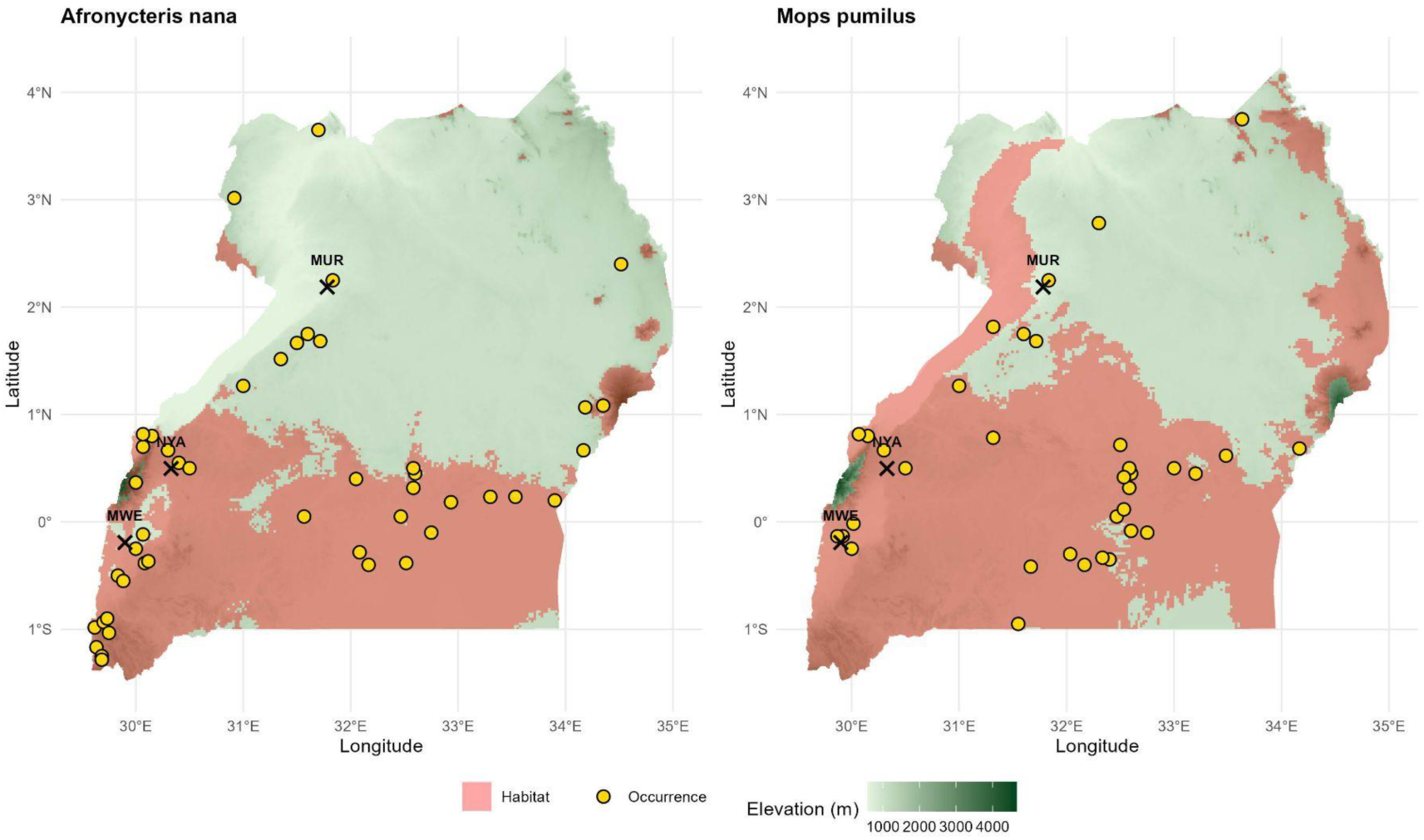
Output of the habitat model of Monadjem et al. (2024) showing the distribution of *A. nana* (left) and of *M. pumilus* (right) at three recording sites (MUR, MWE, NYA; X symbols) at which we observed considerable FM bat activity. Latitude is shown on the y-axis and longitude on the x-axis. The red outlines show suitable habitat for these species, with the gold circles showing areas where specimens have been captured (bat occurrence).

Kaleidoscope Pro identified FM calls at the three sites as consistent with those of the molossid, *Chaerophon (Mops) pumilis* (little free-tailed bat), with match ratios of 0.38 (NYA), 0.47 (MUR), and 0.48 (MWE). The echolocation calls of *M. pumilis* span have low frequency (first harmonic) energy in the frequency range of ∼20-∼40 kHz (South Africa: Fenton et al. 2004, Monadjem et al. 2020; Swaziland: Monadjem et al. 2017). The echolocation calls of *M. condylurus* (Angolan free-tailed bat) have a similar frequency range (∼19-∼37 kHz; Monadjem et al. 2017, 2020). We suggest that both of these molossids may have been present at our recording sites.

## Discussion

Across our six study sites, we recorded high levels of acoustic activity from freely flying bats, with call rates ranging from 43 to 195 calls/min. Some of the variability in call number likely reflects the variable distances of the bats from our recording microphone. Calls were identified in frequency ranges spanning from ∼20 to ∼90 kHz. Due to atmospheric attenuation of very high frequencies (Lawrence and Simmons, 1982) combined with the 192 kHz sampling rate of our recordings, we were unable to detect higher harmonics, thus limiting our ability to identify other high duty-cycle echolocators known to be present in western Uganda (Monadjem et al. 2011, 2020).

Spectral and temporal entropy and entropy index are high at all of our recording sites, with spectral entropy ranging between 0.68 to 0.71, temporal entropy ranging between 0.88 and 0.94, and overall entropy index values ranging from 0.74 to 0.74. Values of these magnitudes confirm the presence of multiple species and an acoustically rich habitat (Sueur et al. 2008).

CF-FM calls are prominent in recordings from MUT, RUB, and RUH, where they were collected at the openings to caves during evening emergence. Parameters of these calls are consistent with those emitted by Rhinolophids; in particular, the second harmonic frequency is consistent with the resting frequency of *R. clivosus,* of which *R. acrotis* may be a lineage, and of *R. alcyone* (Uvizi et al., 2024). Habitat maps suggest that our recording sites are appropriate habitat for *R. acrotis* (Monadjem et al. 2024); however, we were unable to find published literature on the echolocation calls of this species. We cannot rule out the possibility that other CF-FM species with similar echolocation calls were also present in our recordings. Kaleidoscope Pro gave matches of 0.27 at MUT and 0.5 at RUH with *R. blasii*, but our recording sites are not expected to provide good habitat for this species (Monadjem et al. 2024).

FM calls of several different species are prominent at three of our recording sites. The results of the Kaleidoscope Pro analyses indicate that *C. pumilis* is a good match to our recordings. *C. pumilis* has been confirmed to inhabit western Uganda (Monadjem et al. 2011, 2024). From reference to habitat maps and to published literature (Monadjem et al. 2017, 2020, 2024; Taylor-Boyd et al. 2025), we suggest that the vespertilionid *A. nana* was present at one of our recording sites.

Across all recording locations, echolocation calls occurred with temporal overlap yet frequency separation, demonstrating acoustic niche partitioning (Krause 1993; Denzinger and Schnitzler 2013). Visual analyses of spectrograms suggest that multiple individuals of the same species were vocalizing either synchronously or alternately. Although we label all of our recorded signals as echolocation calls, it is possible that some of these calls are used in social interactions and not just in orientation or prey capture.

Although by no means complete, our results provide a glimpse into the bat activity and diversity present in western Uganda. Limitations of our study include the use of only one recording microphone, the sampling rate used for data collection and analysis, sampling only one recording site at each location, and the limited time span of the recordings. Because of permitting restrictions, we were unable to capture bats, so we cannot make definitive species identifications. Passive acoustic monitoring without physical captures is still of use for surveying and for educational purposes (in our case, members of the local populace who assisted us in identifying recording sites were able to “hear” echolocation calls down-sampled by a bat detector to the human audible range). It is important to note that because we intentionally targeted our recordings for time periods and locations of high bat activity, we do not suggest that our reported levels of bat activity and diversity are present across the entire landscape of Uganda. Our recordings highlight some of the bat acoustic diversity in western Uganda, the potential for further exploration, and the need to build call libraries to better understand bat behavior. To contribute to this effort, we have archived our acoustic recordings (see Data Availability) to be of use for the scientific community.

## Author contributions

Field recordings were performed by LNK and RH, with equipment provided by JAS. PRP wrote the MATLAB code. RF, OM, and AMS analyzed the data. LNK performed statistical analyses. LNK, RF, JAS, and AMS wrote and edited the manuscript. All authors approved the final manuscript.

## Acknowledgements

In preparing this publication, we recognize and respectfully acknowledge that some of the research and recordings were conducted on territories traditionally inhabited by the Batwa people. Field recordings were supported by Mugorozi Jackson, Nkusi Gerald, Mugyeni Ezra, the teachers of the Kyamakanda Primary School, and the local residents of Murchison Falls, Mutolere, Mweya, Nyabikere, Rubuguri, and Ruhanga. Renee Choi, Madeleine McLaughlin, Jessica Hooper, and Max Newman assisted with data analysis.

## Data Availability

Audio files recorded for this study are available at the Brown Digital Repository. https://doi.org/10.26300/4s21-8402

